# Noise Stress induces Cardiovascular Metabolic Shifts

**DOI:** 10.1101/2024.04.22.590539

**Authors:** Jair G. Marques, Marin Kuntic, Roopesh Krishnankutty, Giovanny Rodriguez Blanco, Mykyta Malkov, Katie Frenis, Jimi Wills, Engy Shokry, Frederic Li Mow Chee, Cormac T Taylor, Thomas Münzel, Andreas Daiber, Alex von Kriegsheim

## Abstract

Environmental stressors present in the modern world can fundamentally affect humans’ physiology and health. Exposure to stressors like air pollution, heat, and traffic noise has been linked to a pronounced increase in non-communicable diseases. Specifically, aircraft noise has been identified as a risk factor for cardiovascular and metabolic diseases, such as arteriosclerosis, heart failure, stroke, and diabetes. Noise stress leads to neuronal activation with subsequent stress hormone release that ultimately leads to activation of the renin-angiotensin-aldosterone system, increasing inflammation and oxidative stress, dramatically affecting the cardiovascular system. However, despite the epidemiological evidence of a link between noise stress and metabolic dysfunction, the consequences of exposure at the molecular, metabolic level of the cardiovascular system are largely unknown. Here we use a murine model system of aircraft noise exposure to show that noise stress profoundly alters heart metabolism. Within days of exposing animals to aircraft noise, the heart has a reduced potential for utilising fatty-acid beta-oxidation, the tricarboxylic acid cycle, and the electron transport chain for generating ATP. This is compensated by shifting energy production towards glycolysis. Intriguingly, the metabolic shift is reminiscent of what is observed in failing and ischaemic hearts. Our results demonstrate that within a relatively short exposure time, the cardiovascular system undergoes a fundamental metabolic shift that bears the hallmarks of cardiovascular disease.

Overall, aircraft noise induces rapid, detrimental metabolic shifts in the heart, resembling patterns seen in cardiovascular diseases. These findings underscore the urgent need to comprehend the molecular consequences of environmental stressors, paving the way for targeted interventions aiming at mitigating health risks associated with chronic noise exposure in our modern, noisy environments.

## Introduction

Environmental stressors, such as air, noise, and light pollution, have all been linked to non-communicable diseases and are suggested to out-compete the global burden of disease that is caused by genetic predisposition (1, 2). Specifically, the WHO concluded that traffic noise pollution affects cardiovascular disease development and may contribute to metabolic disease (3). Chronic exposure to road, railway, or aircraft noise has been associated with elevated blood pressure, arterial hypertension, stroke, heart failure, and arrhythmia (4, 5). Whereas noise has been identified as a significant cardiovascular risk factor (6), the molecular mechanisms by which noise contributes to the pathogenesis of cardiovascular disease are only partially understood (7).

At the molecular level, noise stress induces the elevation of stress hormones, systemic inflammation, and oxidative stress (8). The latter not only drives atherosclerosis but also potentially fundamentally alters cellular metabolism by restricting oxygen and nutrient supply (9). This metabolic alteration causes arterial stiffness and atherosclerosis, as indicated by the accumulation of lipids and oxidized lipids in atherosclerotic plaques (10, 11).

Lipid oxidation can be driven by an imbalance of oxidants and antioxidants. This further leads to harmful reactive oxygen species (ROS) and reactive nitrogen species (RNS) contributing to inflammation and apoptosis. Furthermore, excessive ROS can progressively damage the mitochondrial respiratory machinery. Together, these processes will lead to the accumulation of harmful by-products and metabolic intermediates prone to oxidation. This initiates a detrimental feedback loop, negatively impacting mitochondrial function and amplifying cardiac structural alterations in a cyclic cascade of “degenerative remodelling” (12). The interconnectedness of these molecular responses underscores the intricate relationship between noise stress and cardiovascular health.

How metabolic pathways are affected at the molecular level by noise stress is still under investigation despite evidence for a strong functional link between noise stress, cardiac disease, and metabolic dysfunction. Here we determined how cardiovascular metabolic networks are dysregulated by acute noise stress in a murine model system to give a novel insight of how an environmental stressor can promote cardiovascular and metabolic dysfunction.

## Results

### Dysregulation of Metabolic Networks in Noise-Stressed Animals

To determine whether any metabolic networks were dysregulated in noise-stressed animals, we analysed a published transcriptomics dataset of murine aortic tissue that was isolated from either control C57BL/6J animals or from a cohort that had been subjected to four days of continuous aircraft noise (13). We used the string database to cluster significantly altered mRNAs and found several highly connected network clusters (Fig. 1a).

**Figure 1:**
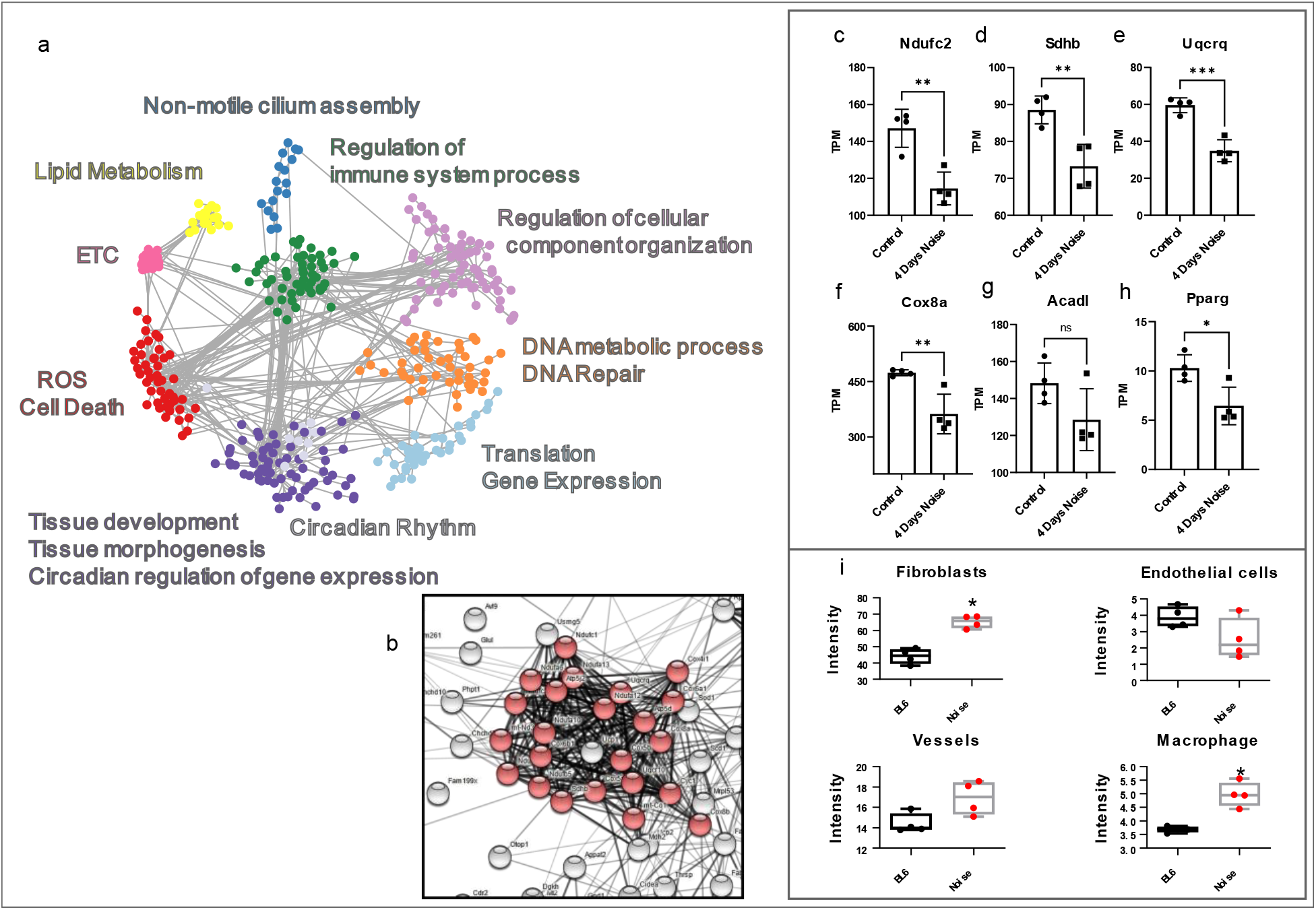
Pathways analysis of Noise-Stressed Aorta identifies dysregulated expression of transcripts of ETC, OXPHOS components and cellular composition. **(a)** Network analysis of mRNA transcripts significantly altered in the aorta from noise-stressed mice. **(b)** Zoom of Mitochondrial ETC cluster **(c-f)** Expression profile of mRNA transcripts from Complex I to IV or the ETC **(g-h)** Expression profile of mRNA transcripts regulating lipid metabolism **(i)** Declustering of RNAseq data identifies changes of cellular composition of aorta upon noise-stress.

Several clusters were mapped to pathways that were expected or were previously reported to be affected by noise, such as circadian rhythm, DNA damage repair, and oxidative stress. In addition, we detected several networks linked to metabolism, chiefly lipid metabolic pathways and the mitochondrial electron transport chain (ETC) (Fig. 1b), which could indicate a noise-induced rearrangement of aortic energy production. Upon closer inspection, we determined that mRNA levels of essential ETC components were significantly downregulated (Fig. 1c-f). In addition, we detected the reduced expression of the Peroxisome proliferator-activated receptor gamma (Pparg) a transcription factor that drives the expression of enzymes essential for the peroxisomal beta-oxidation pathway of fatty acids (Fig. 1g). We also noticed that Acadl, a Pparg-target gene, was significantly downregulated in the aorta of noise-stressed animals (Fig. 1h).

Overall, these data implied that noise stress in animals led to reduced capacity in the aorta to generate energy from fatty acid beta-oxidation via oxidative phosphorylation (OXPHOS), potentially shifting to another means of adenosine triphosphate (ATP) production.

There are two likely explanations for this observation. Either the endothelial and aortic cells have altered their capacity for ATP production or there has been a dramatic shift in the cellular composition of the aorta towards more glycolytic cells, such as immune cells.

### Cellular Composition and Immune Cell Infiltration

To determine if a massive immune cell infiltration could have led to a reduction in the bulk mRNA signal of OXPHOS genes, we used a network-based approach to infer the cellular composition based on bulk mRNA expression. We deconvoluted the bulk mRNA data and inferred the cellular composition of the aorta using ImSig (14) (Fig. 1i). Of the immune cells assayed, only macrophage infiltration was significantly changed by noise stress. We detected around a 30% increase in aorta-resident macrophages, which is in line with published data (13). The endothelial composition of the aorta was unaffected, whereas we detected a significant increase in the fibroblast signature, which could suggest an increase in aortic-resident fibroblasts or fibrosis (15). Conversely, as the overall changes in immune cell infiltration appear to be marginal, we postulate that the observed expression changes of ETC and Pparg-pathways gene transcripts were not only due to the increase in macrophage infiltration.

To determine whether the reduction in the mRNA levels of electron-transfer chain genes was translated into reduced protein levels we reproduced the experiment. Mice were exposed to a recording of aircraft noise on a loop for 9 ± 3 h per day for 3 days, after which the animals were sacrificed. The aorta was isolated and processed for mass spectrometry-based proteomics to quantify the proteome.

### Aorta Proteomic Analysis

We quantified over 4000 protein groups and determined which were statistically significantly regulated upon noise exposure (Fig. 2a). Using enrichment analysis on this subset, we found that proteins relevant to the ETC and respiration were significantly over-represented (Fig. 2 b).

**Figure 2:**
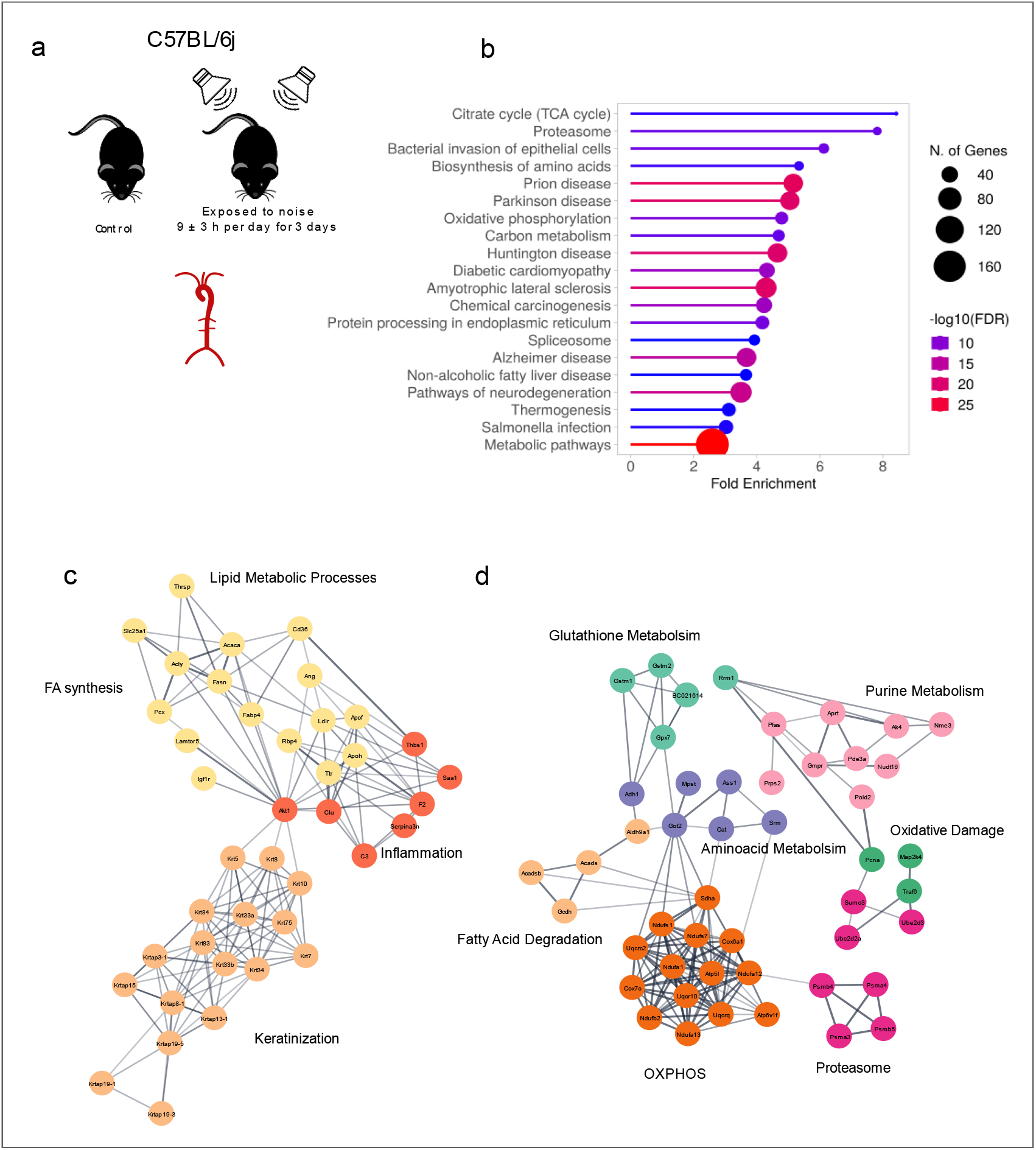
Proteomic Network analysis of Pathways affected in the Aorta by Noise-Stress. **(a)** Gene Ontology Enrichment Analysis of proteins significantly altered by noise stress in murine aortas **(b)** Network analysis of proteins significantly increased by noise-stress in the murine aorta **(c)** Network analysis of proteins significantly decreased by noise-stress in the murine aorta.

Using string network analysis, we discovered that signalling clusters linked to Inflammation, keratinization, and lipid metabolism, particularly fatty acid synthesis, were upregulated (Fig. 2c). Conversely, protein networks linked to OXPHOS, fatty acid degradation were downregulated (Fig. 2d). In addition, clusters that are related to oxidative stress and glutathione metabolism were also enriched. Taken together, these data imply that noise stress could reduce the metabolic capacity of the aorta to catabolise fatty acids by beta-oxidation and generate ATP via the ETC. We also detect inflammation and oxidative stress signs, which are previously identified hallmarks of noise stress in the cardiovascular system (13). Broadly, the changes in the proteome are consistent with what we observed in the transcriptome, both of which suggest a major rewiring of the aortic ATP-producing capacity

### Cardiac Impact of Noise Stress

To ascertain if this noise-induced shift in metabolic potential was mirrored across the cardiovascular system, we measured the cardiac proteome of noise-stressed animals to identify noise-regulated proteins and networks. As done above, we processed the hearts of stressed animals using a recording of aircraft noise on a loop (9 ± 3 h per day for 3 days).

Enrichment analysis of significantly regulated proteins revealed that proteins associated with fatty acid metabolism and OXPHOS were significantly enriched in the regulated dataset (Fig. 3a). In addition, proteins associated with cardiomyopathy were also enriched.

**Figure 3:**
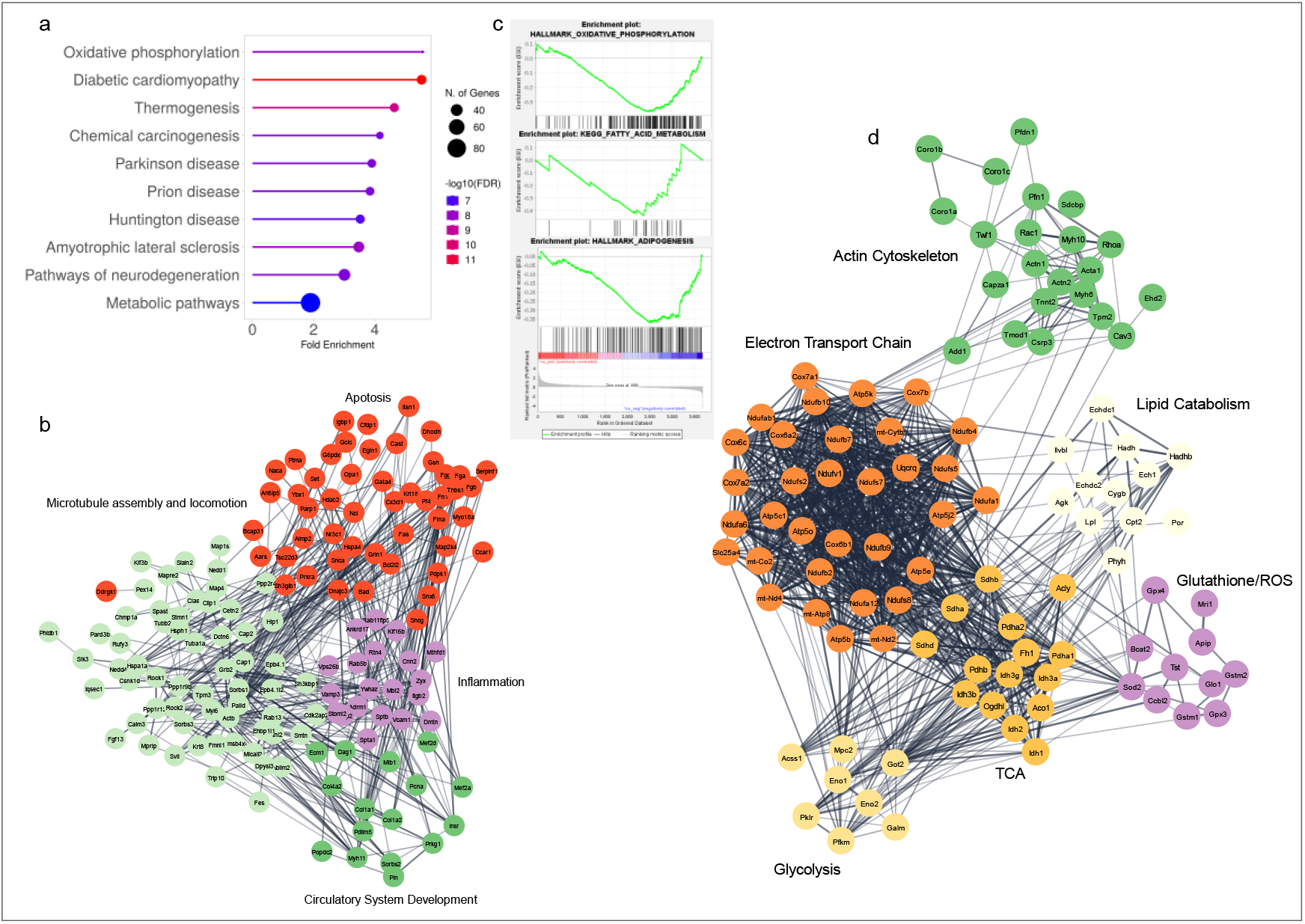
Proteomic Network analysis of Pathways affected by by Noise-Stress in the Heart. **(a)** Gene Ontology Enrichment Analysis of proteins significantly altered by noise stress in the murine heart **(b)** Network analysis of proteins significantly increased by noise-stress in the murine heart **(c)** Three significant Hallmarks of pre-ranked by ratio Noise/Control GSEA analysis of heart proteome **(d)** Network analysis of proteins significantly decreased by noise-stress in the murine heart.

Network analysis of the proteins elevated in noise-stressed hearts revealed four interconnected networks composed of an inflammatory and an apoptotic signature (Fig. 3b). We also detected two clusters linked to microtubule dynamics and circulatory systems development. In contrast, when looking at downregulated proteins, we found a tightly connected cluster comprising the ETC, TCA cycle, glycolysis, and lipid catabolism (Fig. 3c). We further identified a cluster associated with ROS and the actin cytoskeleton. Also, we found several proteins that sit at the crucial interphase of the interconnected networks. These proteins include the succinate dehydrogenase complex, part of the TCA cycle and ETC, or GPX4, a protein glutathione peroxidase essential for detoxifying peroxidised lipids.

### Analysis of ETC and Lipid Metabolism in the Heart

The dysregulated networks were considerably larger and more interconnected in the heart when compared to what we observed in the aorta. Therefore, we decided to investigate the nature of the regulated proteins in more detail. We initially plotted all ETC proteins that were found to be significantly regulated on a volcano plot (Fig. 4a). It became apparent that regulation was not uniform across all four complexes. The significantly regulated Complex I proteins were all downregulated (Fig. 4b). Also, three of the found succinate dehydrogenase (SDH) subunits of Complex II were significantly downregulated, and the fourth was trending downwards (Fig. 4c). The picture was more mixed for Complex III and IV subunits where we detected up and downregulated proteins (Fig. 4a). Next, we looked at the lipid metabolic/catabolic/transport pathways (Fig. 4d). We found that core proteins of several lipid catabolic pathways were downregulated (Fig. 4e), whereas proteins involved in lipid shuttling were overexpressed in noise-stressed animals (Fig. 4f).

**Figure 4:**
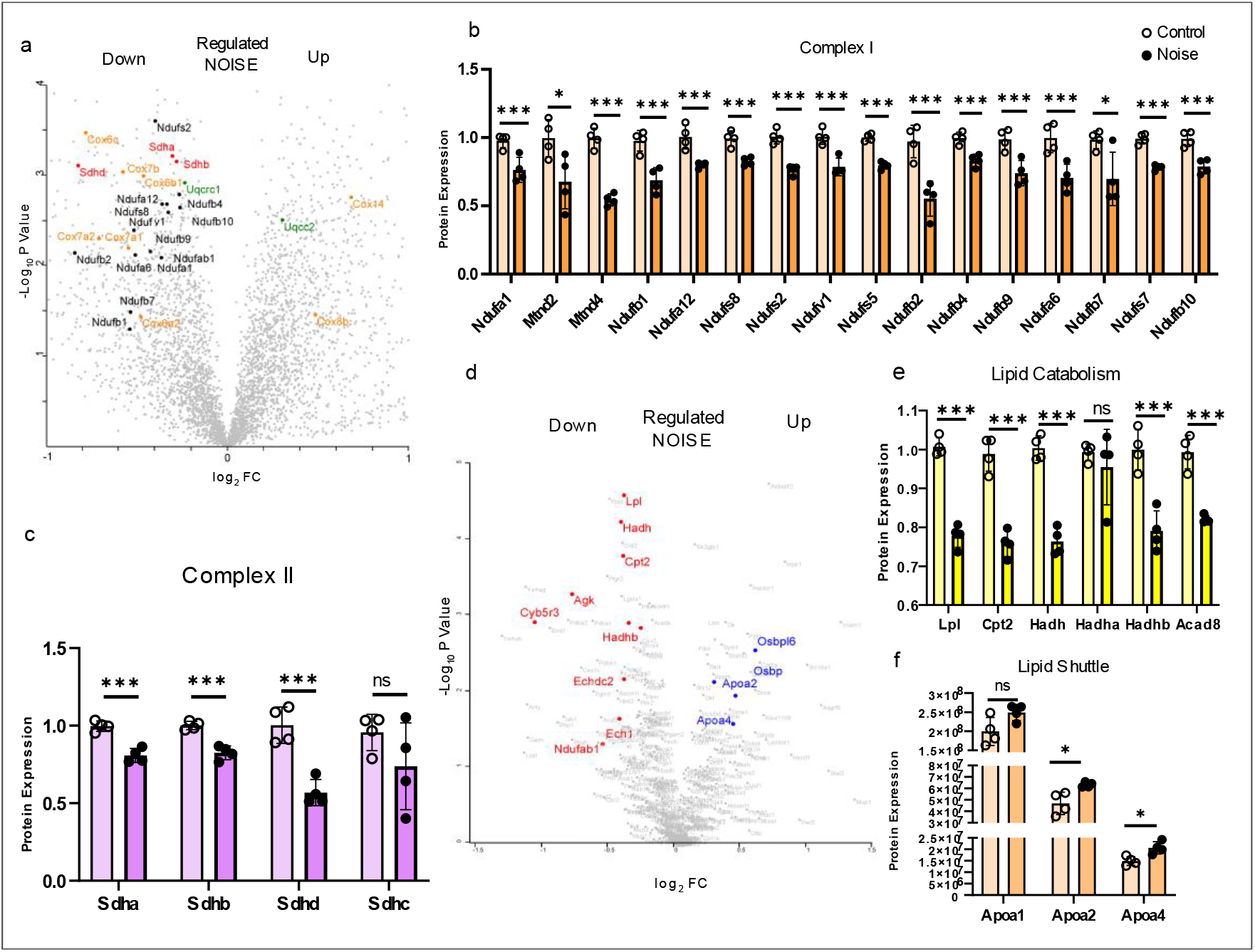
Disruption of ETC and lipid metabolism in the Noise-stressed Heart. **(a)** Volcano plot of Heart Proteome depicting significantly altered ETC proteins with Complex I to IV as black, red, green and yellow, respectively **(b)** Expression changes in Complex I proteins with empty circles set as Control and full circles as Noise stress **(c)** Expression changes in Complex II proteins with empty circles set as Control and full circles as Noise stress **(d)** Volcano plot of Heart Proteome depicting significantly altered lipid metabolism proteins with proteins involved in lipid catabolism as red and lipid transport in blue **(e)** Expression changes of lipid catabolism proteins with empty circles set as Control and full circles as Noise stress **(f)** Expression changes of lipid transport proteins with empty circles set as Control and full circles as Noise stress

Together, these data suggest that noise stress is likely to affect the capacity of major metabolic pathways to generate ATP. Given the plasticity of cardiac metabolism, we cannot conclude if changes in expression levels result in an actual shift in metabolism. Although suggestive, the data are also partially contradictory. We detected the simultaneous downregulation of rate-limiting enzymes involved in glycolysis, beta-oxidation, the TCA cycle and the ETC, which would suggest a reduction in the overall capacity of the heart to generate ATP, but we have no indication if and how the major means and pathways for generating energy are affected by noise stress. To determine whether noise stress affects catabolic and metabolic pathways, we isolated hearts from animals stressed using aircraft noise Fig. 5a and used a biphasic extraction protocol (16) to isolate hydrophilic and hydrophobic metabolites.

**Figure 5:**
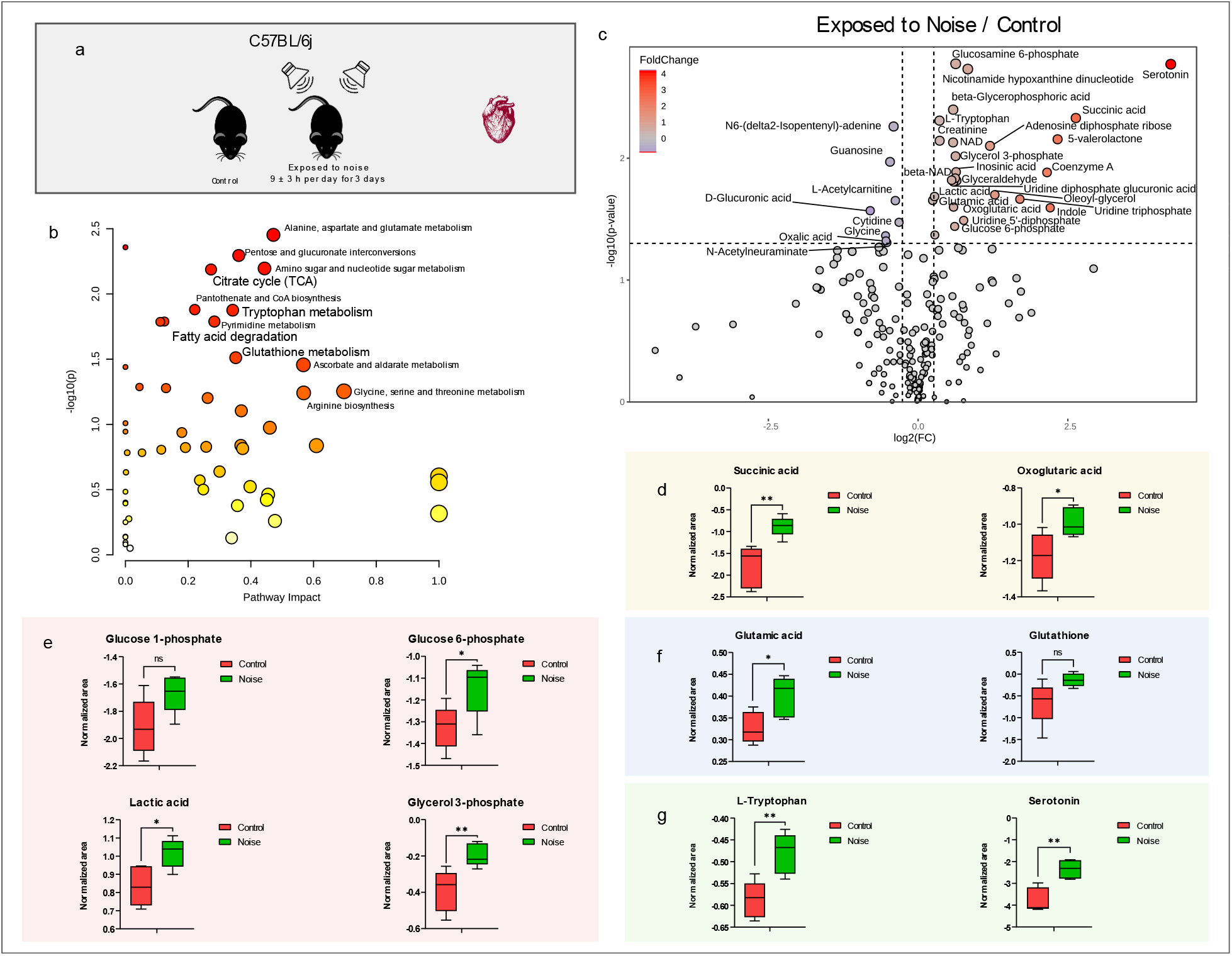
Noise-stress-induced Glycolysis and impeded lipolysis. a. Experimental setup: Hearts isolated from animals exposed to aircraft noise stress (b) Metabolic pathway perturbations: Significant changes identified in key metabolic pathways in response to noise stress (c) Volcano plot: Visualization of individual metabolites, indicating their impact on pathways (d) Tricarboxylic acid (TCA) cycle: Elevated levels of metabolites oxoglutaric acid and succinate (e) Glycolysis and Lipogenesis: Increased levels of glucose 6-phosphate, non-significant increase in glucose 1-phosphate, elevated lactic acid, and Glycerol-3-phosphate, indicating a shift in ATP production towards glycolytic pathways (f) Oxidative stress response: Alterations in glutathione metabolism are highlighted by elevated glutamic acid levels and a non-significant increase in glutathione, indicating oxidative stress (g) Tryptophan and Serotonin: Increased levels suggest an adaptive response to counteract oxidative stress induced by noise exposure. Serotonin significantly increased, indicating an association with cardiac function.

### Metabolomic Profiling Unveils Shifts in Metabolic Pathways

We identified significant perturbations in key metabolic pathways in response to noise stress (Fig. 5b). Observing the metabolites individually on a volcano plot (Fig. 5c) we can visualize how the pathways are impacted. The TCA cycle exhibited increased levels of metabolites such as oxoglutarate and succinate (Fig. 5d). lactic acid, and Glycerol-3-phosphate, indicating a shift in ATP production towards glycolytic pathways (f) Oxidative stress response: Alterations in glutathione metabolism are highlighted by elevated glutamic acid levels and a non-significant increase in glutathione, indicating oxidative stress (g) Tryptophan and Serotonin: Increased levels suggest an adaptive response to counteract oxidative stress induced by noise exposure. Serotonin significantly increased, indicating an association with cardiac function.

Succinate was amongst the most significantly increased metabolites and was induced over 5-fold compared to the control. The elevated succinate levels align with the observed downregulation of SDH complex proteins, indicating a potential blockage in the TCA cycle at the level of succinate. Interestingly, the abundance of SDH proteins was reduced in noise-stressed hearts (Fig. 4c) only by a maximum of a 2-fold reduction, making the over 5-fold increase in succinate notable.

Glucose 6-phosphate was significantly increased in noise-stressed animals, and a non-significant increase of glucose 1-phosphate (p = 0.057) was detected. Furthermore, lactic acid, the product of the pathway most indicative of oxygen-independent ATP production, and glycerol-3-phosphate, a metabolite linking glycolysis to the mitochondria activity via its shuttle, significantly increased (Fig. 5e). Overall, these data demonstrate a shift of ATP production via the TCA cycle and ETC towards glycolytic ATP production.

Glucose 6-phosphate was significantly increased in noise-stressed animals, and a non-significant increase of glucose 1-phosphate (p = 0.057) was detected. Furthermore, lactic acid, the product of the pathway most indicative of oxygen-independent ATP production, and glycerol-3-phosphate, a metabolite linking glycolysis to lipogenesis, significantly increased Fig. 5e. Overall, these data demonstrate a shift of ATP production via the TCA cycle and ETC towards glycolytic ATP production.

The decrease in L-acetylcarnitine (Fig. 5c) indicates a compromised fatty acid transport for beta-oxidation. Concurrently, alterations in glutathione metabolism, highlighted by the significant elevation in glutamic acid levels and a non-significant elevation in glutathione (p = 0.056) (Fig. 5f) is a hallmark of the oxidative stress response. These findings indicate a multifaceted impact of noise stress on cardiovascular metabolism, influencing energy production, fatty acid metabolism, and antioxidant defences.

### Complex Impact on Cardiovascular Metabolism

Beyond the shift in energy metabolism, we detected several other pathways implicated in the response to noise stress. Amino acids metabolism pathways (alanine, aspartate, and glutamate metabolism; Glycine, serine, and threonine metabolism; Arginine biosynthesis) were significantly impacted, as well as amino sugar and nucleotide sugar metabolism (Fig. 5b). However, the amino acid tryptophan was the most prominent amino acid detected in the univariate analysis. On close observation, levels of tryptophan and serotonin were significantly increased (Fig. 5g). Tryptophan regulates the immune response and antioxidant defence mechanism. An increase suggests an adaptive response to counteract the oxidative stress induced by noise exposure. The most significantly increased metabolite detected, serotonin, is known for its association with cardiac function (17). The interplay between tryptophan and serotonin, amidst the identified metabolic dysregulations, implies a complex network of biochemical alterations.

Next, we analysed the lipid-containing hydrophobic fractions. Using a mass spectrometry-based lipidomic approach, we identified and quantified cardiac lipids from both groups. We used differential analysis to determine lipids whose abundance was statistically significantly altered when comparing the control and noise-stressed groups (Fig. 6a). The most prominent lipid class to be decreased in the noise-stressed group was acylcarnitines (AC). AC functions as carriers for free Fatty Acids (FA) into the mitochondria, and deregulation can indicate a decreased FA beta-oxidation pathway. In contrast, we detected increased phosphatidylcholine (PC) species containing polyunsaturated fatty acids. The observed increase in polyunsaturated PCs suggests a dynamic response to noise-induced stress, potentially reflecting adaptations in membrane composition to mitigate the effects of oxidative stress and/or inflammation.

**Figure 6:**
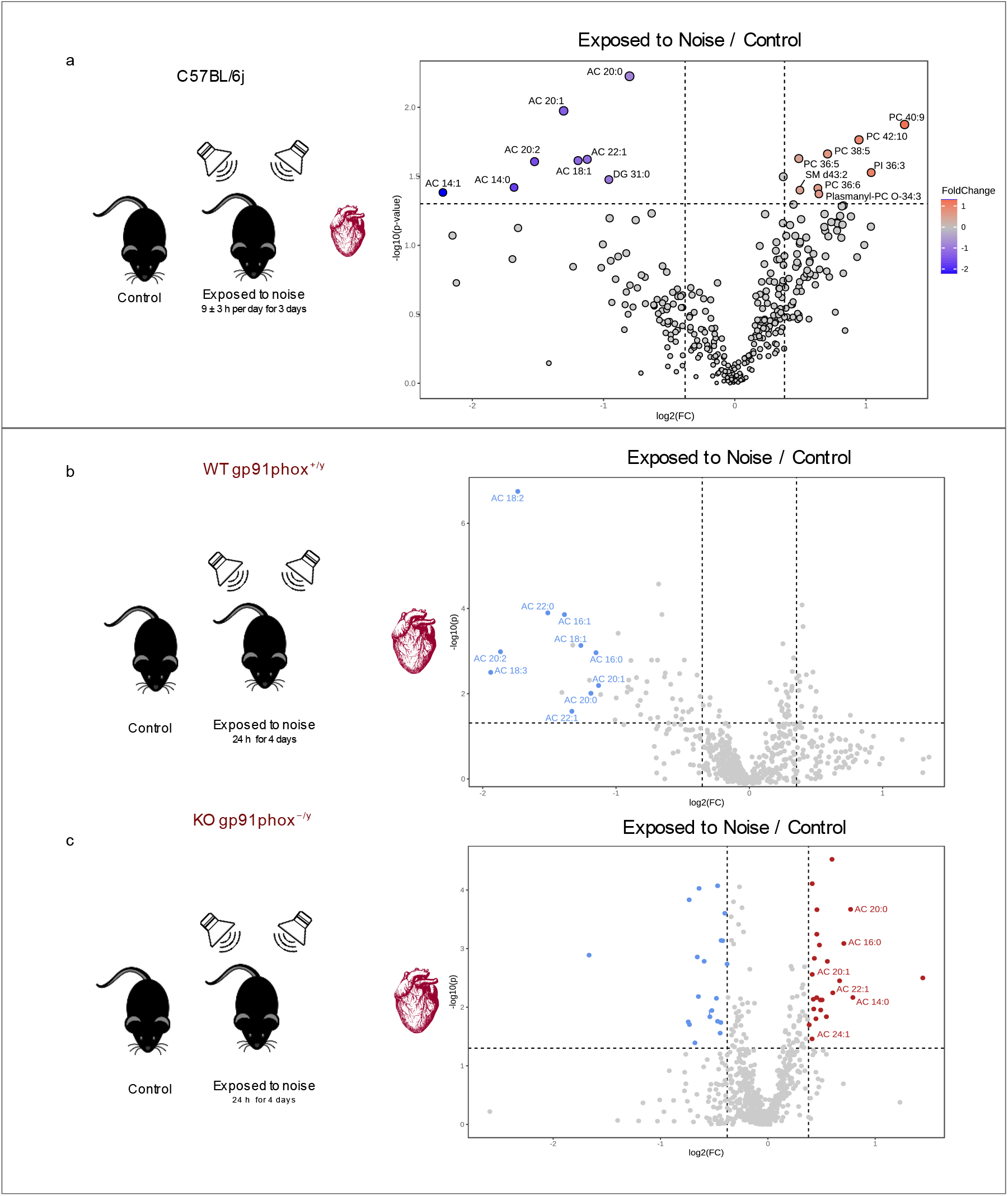
impact of noise stress on cardiac lipid metabolism. (a) Experimental setup and volcano plot determine statistically significant alterations in lipid abundance (p<0.05) (b) Wild-type (WT) animals: Lipidomics experiment results showing an overall decrease in acylcarnitines (ACs) in response to aircraft noise stress, suggesting a reduction in the heart’s capacity for catabolizing fatty acids (c) GP91phox knockout (KO) animals: Lipidomics experiment results in GP91phox KO animals, indicating a significant increase (p<0.05) in ACs even under aircraft noise stress. This suggests a potential rescue of the heart’s capacity for AC-driven fatty acid import and beta-oxidation in GP91phox KO animals.

In summary, the lipidome and metabolome data provide a clear picture, of how in the heart of noise-stressed animals, the means of generating ATP shift away from beta-oxidation and the TCA cycle towards the anaerobic glycolytic pathway with lactate as the final product.

The metabolome data is backed up by our detection of a marked decrease in enzymes essential for lipolysis, including the rate-limiting carnitine palmitoyltransferase (CPT2) (Fig. 4e), as well as the reduction of proteins essential for beta-oxidation, the ETC and the TCA cycle (Fig. 3d). Our finding that glycolytic enzymes were also downregulated, is at first glance inconsistent with the observed increase in glucose metabolism and glycolytic activity. However, flux through metabolic networks can be controlled by altering the activity of intercellular enzymes but can also be increased by potentiating the import of substrates and export of products, glucose, and lactate, respectively, in the case of glycolysis.

We reinvestigated the proteomic data and found that GLUT4, the major cardiac high-affinity glucose importer, was overexpressed in the heart of noise-stressed animals, as was the monocarboxylate transporter 1 (MCT1), a lactate exporter (Supplementary Fig. 1a). This suggests an increased capacity to import the substrate and export the pathway’s product, thereby increasing the capability to generate ATP via glycolysis. Metabolites have been shown to regulate the enzymatic activity via the substrate, product or allosteric activation or inhibition. TCA metabolites, including fumarate, succinate and 2-hydroxy-glutarate inhibit the activity of 2-oxoglutarate-dependent oxygenases (13), including the HIF-regulating EGLN enzyme family that acts as oxygen and metabolite sensors. We were curious to determine whether the observed accumulation of succinate might affect hypoxia signalling by inhibiting EGLN hydroxylase and would therefore indirectly contribute toward the shift towards glycolysis.

We looked at several proteins as markers of hypoxia (Supplementary Fig. 1 b) and found that the negative feedback target of HIF-signalling EGLN1 was indeed induced in noise-stressed hearts, as were carbonic anhydrase 1 and 2 (CA1 & 2), whereas pyruvate dehydrogenase kinase 1 (PDK1), another target of HIF, was unaffected. Overall, these data suggest that the heart tissue is adapting to increased lactate-driven acidification by expressing CAs and the induction of a restricted set of HIF-target proteins. Nevertheless, HIF signalling is a priori a transcription program, therefore, to determine whether we could detect a transcriptional signature consistent with activation of HIF, we interrogated a previously published transcriptomics data set of noise-stressed murine hearts (18). We performed an enrichment analysis on transcripts significantly increased in noise-stressed hearts and, amongst others, indeed found that “HIF-1 signalling pathway” was one of the top enriched signatures (Supplementary Fig. 2a-b), suggesting that HIF activity is at least partially augmented by noise-stress in the heart muscle.

### GP91phox Knock-Out Rescues Phenotype

Next, we sought to determine whether oxidative stress may be the driver of the reduced capacity of the heart to catabolise fatty acids for ATP production as implied by the reduction of AC. Furthermore, we wanted to ascertain if HIF protein expression was induced and whether this was driven by oxidative stress. To test both hypotheses, we analysed archival samples from wild-type (WT) and animals where NOX2 (GP91phox) was knocked out (KO). The GP91phox KO has been previously shown to blunt ROS induction following chronic hypoxia (19) and rescuing some of the phenotypes associated with noise-stress (20, 21). To determine the shift from beta-oxidation to anaerobic glycolysis, we extracted lipids from WT and GP91phox KO that were or were not stressed by aircraft noise using an equivalent exposure protocol as described in (13). The results of the lipidomics experiment of the wild type animals (Fig 6b) showed an overall decrease in AC’s, reproducing the observation obtained in our previous experiment. Remarkably, in the GP91phox KO animals we observed a subtle increase of AC’s (Fig 6c), suggesting that the heart’s potential for AC-driven fatty acid import and beta-oxidation was rescued in the KO animals.

To determine if HIF was induced at the protein level, we performed immunohistochemical staining of heart tissue, specifically for detecting Hypoxia-Inducible Factor 1-alpha (HIF-1α) (Supplementary Fig. 3). However, despite being able to detect HIF1/2 in positive control, we were unable to detect HIF in the heart muscle and can therefore not conclude whether HIF plays a role in the noise stress response or can indeed be rescued by GP91phox KO.

## Discussion

There is a close association between noise exposure and risk of cardiovascular events, like hypertension, arrhythmias, myocardial infarction, and stroke in large cohorts (22-27). Other large cohort studies found a significant association between noise exposure and energy metabolism changes, like the risk of developing metabolic syndrome and insulin resistance (28, 29). These metabolic changes by noise were also confirmed in animal studies (30, 31). In addition to metabolic changes, animal studies also showed that noise was associated with mitochondrial dysfunction and structural alterations together with disturbance of respiratory chain activity (32-34). Noise promotes a stress response, resulting in the release of hormones like cortisol, which can, in an acute setting, promote lipolysis, or in a chronic setting, reduce lipolysis (35), although some studies observed increased lipolysis even in a chronic stress setting (36). Disease burden by transportation noise and the intricate changes to oxidative stress and metabolic regulations were recently reviewed (37). Whereas the impact of noise exposure on metabolic processes is evident, the understanding of the underlying pathomechanisms is incomplete. This was the starting point and rationale for the present study. Under physiological conditions, the cardiovascular system derives 70-95% of its ATP production by catabolising lipids through the beta-oxidation pathway (38, 39). The healthy heart is metabolically very flexible, it has overcapacity in the catabolic pathways. It can rapidly alter energy production from one fuel source to another, and anaerobic glycolysis can generate ATP more quickly than beta-oxidation. In situations of sudden and intense energy demands, such as during acute stress or ischemic events, the heart may activate glycolysis to rapidly produce ATP (40). Here we show that noise stress induces the readjustment of cardiac metabolism by shifting from fatty acids beta-oxidation to anaerobic glycolysis as a source of ATP production. This metabolic phenotype is similar to that observed In advanced heart failure, with reduced fatty acid oxidation, increased glycolysis and glucose catabolism, decreased respiratory chain activity, and impaired mitochondrial oxidative flux (39). In our study, succinate was one of the most significant metabolites upregulated by noise stress, an intermediate of the TCA cycle that is generally quickly converted to fumarate by the SDH/Complex II of the ETC/TCA cycle. In the stressed heart, succinate accumulates 5-fold over baseline. The question is why this is occurring. We detect a reduction of the entire SDH complex at the proteome level, but the reduction ranges from 10 to 50%, possibly too marginal to explain the accumulation of succinate to the extent we have observed. One possible explanation is that a sizable proportion of the SDH complex is inactivated, possibly by the presence of ROS. We have detected markers of oxidative stress and ROS in the proteome and metabolome, and it is one of the postulated hallmarks in the stressed cardiovascular system (41). Further work should establish if ROS contributes or may even be the initiating event of succinate accumulation and downregulation of the TCA cycle, and ETC. Nevertheless, as we did not uncover any evidence that TCA cycle flux is enhanced, the accumulation of a TCA intermediate succinate, strongly suggests that the enzyme downstream has become a chokepoint. The main consequence of it is the apparent loss of metabolic plasticity. The presence of such an eye of a needle indicates that the TCA cycle is at full capacity and the cells are forced to shift towards cytosolic anaerobic glycolysis for ATP synthesis to keep up with the energy demand upon stress. Secondly, the accumulation of metabolites will not only impinge on one metabolic reaction, but it will also lead to the backing up of the pathway. Something we have observed in the accumulation of 2-oxoglutaric acid (2OG), the metabolite preceding succinate in the TCA cycle. As 2OG is a key substrate for enzymes involved in epigenetic regulation, such as the Jumonji C domain-containing histone demethylases (KDMs), changes in 2OG levels can affect histone demethylation and, consequently, gene expression, linking cellular metabolism with epigenetic regulation. Finally, succinate is the product of the protein hydroxylation reaction elicited by 2OG-dependent hydroxylases and is a potent inhibitor of the reaction due to product inhibition.

We further observed that noise stress regulates lipid metabolism at the protein level. Significant downregulation of Lipoprotein lipase (LPL), a key enzyme involved in the hydrolysis of triglycerides (TGs) from circulating lipoproteins, points to an inhibition of lipid oxidation when observed concomitantly with decreased expression of carnitine palmitoyl transferase 2 (CPT2), the last step importing fatty acid into the mitochondria for beta-oxidation. The result is the observed decrease in acyl-carnitines, closing the picture of the diminished beta-oxidation process. The upregulation of apoA-1, apoA-2, and apoA-4 may contribute to the enhanced transport of lipids, potentially as a response to maintain lipid homeostasis under stress conditions. AMPK signalling pathways were also highly regulated in the noise-exposed mice (Supplementary Fig 2 A). In our previous study, we showed that activation of AMPK by pharmacological treatment, exercise, or fasting, had a beneficial effect on aircraft noise-induced vascular dysfunction and oxidative stress, and protective effects were abrogated in AMPK-deficient mice (42). AMPK pathway plays an important role in energy homeostasis, and modulation of this pathway could explain the lower utilisation of fatty acid oxidation in favour of glycolysis, which also aligns with the finding that AC-driven fatty acid import and beta-oxidation were rescued in the GP91phox KO animals.

The observed upregulation of glucose/lactate transporters in the heart suggests an adaptive response, potentially driven by increased levels of succinate or the choking of the TCA cycle. However, the lack of a clear HIF signature and the rescue of the phenotype in GP91phox knockout animals indicates a complex interplay of metabolic alterations, emphasizing the multifaceted impact of noise stress on cardiovascular metabolism and signalling pathways. Moreover, we detected heightened levels of OPA1 (Supplementary Fig 4) suggesting that the OXPHOS system is operating at its peak. OPA1, a key regulator of inner mitochondrial membrane (IMM) fusion, contributes to an interconnected mitochondrial network, optimizing oxidative capacity. Its involvement in cristae remodelling reinforces its role in OXPHOS regulation. The increase in OPA1 levels reflects both the energetic demands on mitochondria and a response to cellular stress. OPA1-mediated mitochondrial fusion serves as a mechanism to mix contents, rescue damaged mitochondria, and potentially alleviate stress-induced challenges (43). Although HIF expression changes by noise were not detected at the protein level, the transcriptomic activation of this signalling pathway suggests a certain reaction to a hypoxia-like state. Our previous study in a mouse model of myocardial infarction (MI) showed that noise exposure before MI substantially amplifies subsequent cardiovascular inflammation and oxidative stress and worsens ischaemic heart failure (44). The induction of a hypoxia-like state by noise exposure could explain the additive effect on MI, although it is not clear whether chronic noise exposure would lead to ischaemic preconditioning (45).

Another notable observation in the metabolomics data, was the remarkable elevation of serotonin (5-HT) concentration in the noise-stressed group. When considered alongside the significant rise in tryptophan, a precursor in 5-HT synthesis, it suggests a localized upregulation of 5-HT within the cardiac tissue. This suggests that the increased tryptophan and 5-HT levels may intricately influence metabolic pathways, contributing to the adaptive responses observed in the mice subjected to noise-induced stress. Previous research has reported the presence of 5-HT in mammalian hearts and cardiomyocytes, underscoring its significance in cardiac function (17, 43, 46, 47). When oxidized in mitochondria, 5-HT generates free radicals, leading to intracellular effects and the potential induction of apoptosis and necrosis (48). Interestingly, Sola-Penna et al. described how increasing concentrations of 5-HT promoted glucose consumption and lactate production by MCF-7 cells via receptor-dependent fashion (49), a similar phenotype observed in our results. Furthermore, 5-HT can promote protein serotonylation, a receptor-independent covalent modification catalysed by transglutaminases, impacting the function of proteins within cardiomyocytes. This includes connections to fibrinogen, small G-proteins, and even histones, suggesting a direct influence on cardiac gene transcription without receptor involvement (50). The receptor-independent actions of 5-HT and its positive inotropic effects observed in various mammalian species emphasize its multifaceted role in cardiac physiology and its potential influence on signal transduction and gene expression (51).

One limitation of the present study is the translational potential of the observations to the human cardiovascular system. The murine audio-sensory system is more sensitive, and the animals could be more susceptible to patterns of aircraft noise, potentiating the effects culminating in an over-stressed state. In addition, although we observed no adaption of the mice to chronic noise exposure for 4 weeks PMID: 35174211, major differences between mice and men regarding stress responses and resilience mechanisms may be assumed. However, many of the metabolites and proteins we have detected as increased in the murine heart are known biomarkers of cardiovascular disease in humans, suggesting a reasonable level of conservation across the species. The present study’s findings shed light on the mechanism by which noise interferes with cardiovascular energy metabolism and provide molecular insight into the observations made by previous epidemiological studies. Shift from fatty acid beta-oxidation to glycolysis is not only a known mechanism associated with cardiovascular disease but a potential treatment target for future noise-mitigation strategies. Further studies in humans are needed to fully elucidate the complex interplay between noise and the cardiovascular energy metabolism.

