## Supplementary figures and images for "Noise Stress induces Cardiovascular Metabolic Shifts"

### Supplementary Fig. 1

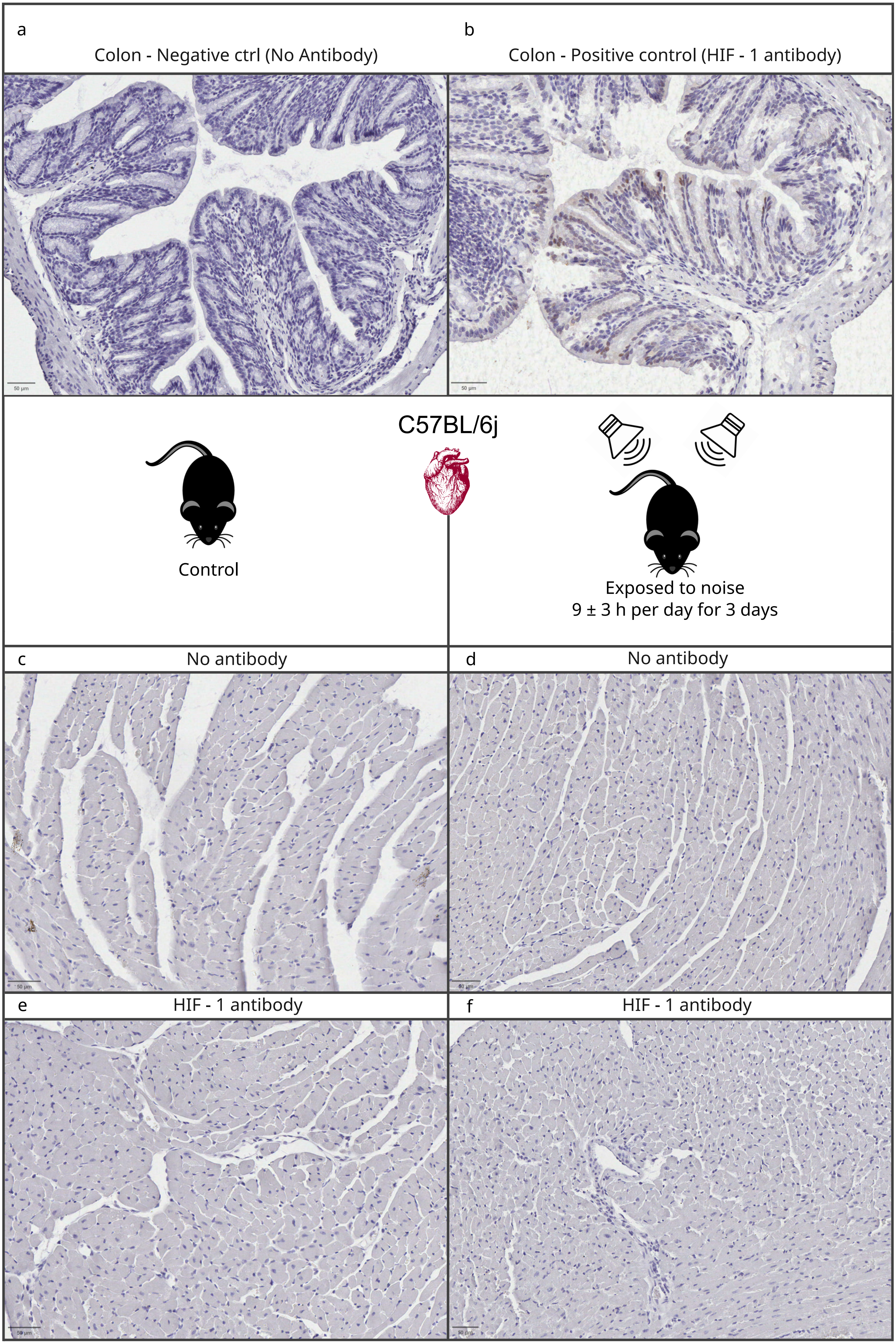

### Supplementary Fig. 1

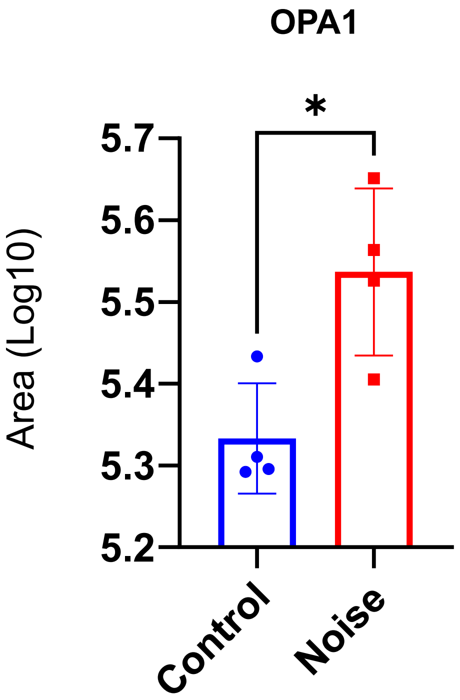

### Supplementary Fig. 3

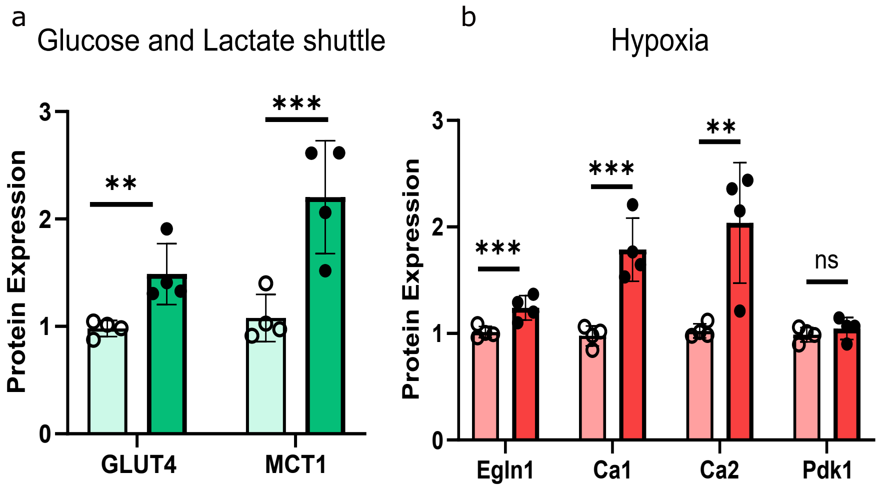

### Supplementary Fig. 4

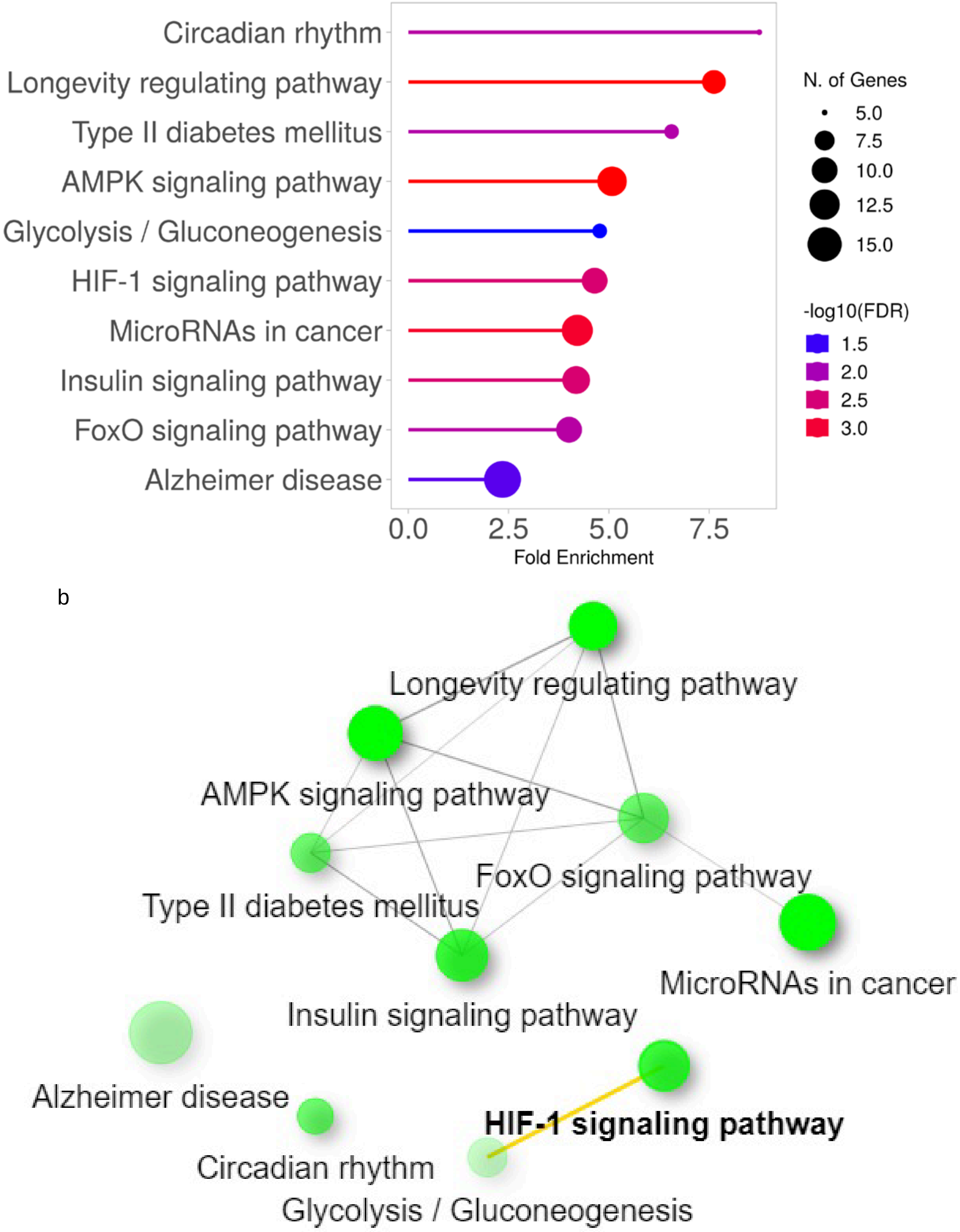
